# Molecular Crowding-Driven Nucleosome Interactions Revealed Through Single-Molecule Optical Tweezers

**DOI:** 10.64898/2026.01.20.700467

**Authors:** Tomoko Sunami, Amarjeet Kumar, Shoko Sato, Yuko Hara, Hitoshi Kurumizaka, Hidetoshi Kono

## Abstract

Molecular crowding causes the compaction of chromatin fibers, contributing to the formation of the nuclear architecture. However, the molecular mechanism of compaction under crowded conditions is not yet fully understood. In this study, we employed the single-molecule optical tweezer method to investigate the effect of molecular crowding on chromatin structure. Force-extension experiments on a 12-mer polynucleosome in the presence of different sizes and concentrations of polyethylene glycol (PEG) as a crowding agent showed that at low concentrations of low-molecular-weight (MW) PEG, the compaction of the polynucleosome was not significant. In this respect, nucleosomes predominantly remained separated, while DNA-histone interactions within individual nucleosomes were slightly stabilized. In contrast, high concentrations of high-MW PEG significantly promote internucleosomal interactions, leading to highly compact polynucleosome conformations. Under these conditions, approximately 30 pN of force was required to disrupt the internucleosomal interactions and release DNA; this force was 36% higher than that required for DNA unwrapping in the absence of PEG. These findings suggest that molecular crowding impacts cellular processes by mechanically regulating chromatin accessibility for regulatory proteins and the passage of motor molecules such as RNA polymerase.

**Significance Statement:** Chromatin condensation is closely related to biological processes such as transcription and replication. Molecular crowding has recently attracted attention as a factor regulating chromatin condensation. In this study, we used the optical tweezer method to analyze the molecular mechanisms underlying chromatin condensation. We found that high-molecular weight and high-concentration crowders (polyethylene glycol) induced significant compaction, which involved internucleosomal interactions that markedly reduced DNA accessibility. Our results suggest that molecular crowding not only alters the condensation state, but also mechanically regulates chromatin accessibility.

## Introduction

Eukaryotic chromatin is composed of long fibers comprising nucleosomes, which are protein-DNA complexes consisting of histone proteins H2A, H2B, H3, and H4, and approximately 147 bp of DNA wrapped around the histone proteins. Chromatin relaxation/condensation is closely associated with gene transcription and replication (1, 2). In cells, the condensation state of chromatin is regulated by the association of nuclear proteins such as heterochromatin protein 1 (HP1), cohesion, and condensin complexes (1, 3–5). Epigenetic markers, particularly post-translational modifications of histone tails, contribute to condensation (6, 7), and environmental factors, such as the concentration of divalent cations (*e*.*g*., Mg^2+^), also play a significant role (8).

Molecular crowding in cells has recently attracted attention as an environmental factor that can influence the chromatin structure. A significant fraction (typically 20–30%) of the total cellular volume is occupied by macromolecules (9), and more than 10% of the nucleoplasm volume also consists of macromolecules (10, 11). The concentration of macromolecules increases further in the nuclear compartments, such as the nucleoli, Cajal bodies, and speckles (10), and under such crowded conditions, the excluded volume effect generates attractive forces between the macromolecules (*i*.*e*., depletion forces) (12).

Several experimental approaches have shown that chromatin compaction can be induced by altering the molecular environment. When HeLa and MCF7 cells were treated with 40–640 mM sucrose, osmotic dehydration increased the intracellular molecular concentration, leading to chromatin compaction and the formation of a new nuclear compartment between the peripheral chromatin and nuclear lamina (13, 14). Similar chromatin compaction was observed in permeabilized cells treated with polyvinyl pyrrolidone or dextran (13–15). More recently, orientation-independent DIC imaging has revealed that the molecular density in the chromosome milieu gradually increases during mitosis from prophase to anaphase, contributing to chromosome condensation (11). In addition, fluorescence microscopy showed that polynucleosomes reconstituted on 166 kbp T4 DNA undergo compaction in solutions containing 10 kDa poly(ethylene glycol) (PEG), monovalent (100 mM Na^+^ and K^+^), and 4 mM divalent (Mg^2+^) cations (16). Bovine serum albumin (BSA) also induces polynucleosome compaction (17). However, while these microscopic analyses have provided structural visualization, they have provided limited insights into the associated molecular mechanisms, such as the specific modes of nucleosome-nucleosome interactions and their thermodynamic stability.

Optical and magnetic tweezers have emerged as powerful tools for analyzing DNA-DNA and protein-DNA interactions at the single-molecule level (18). They can be used to manipulate a single molecule of DNA or protein-DNA complexes and measure the forces acting on them, providing quantitative information about their conformational stability. They have been applied to study DNA condensation induced by ethanol and cations (19), loop formation of DNA mediated by *E. coli* nucleoid proteins, such as HU, IHF, and H-NS (20–23), and nucleosome bridging facilitated by the eukaryotic transcription repressor protein PRC2 (24).

In this study, we used optical tweezers to investigate chromatin structure and stability under molecular crowding conditions. We performed force-extension experiments on 12-mer polynucleosomes in the presence of PEG with varying molecular weights (MWs) and concentrations. An analysis of the force-distance curves revealed the extent of chromatin compaction, the nature of internucleosomal interactions, and the forces required to disrupt these interactions. Our results show that high concentrations of high-MW PEG induce pronounced chromatin compaction, accompanied by stable internucleosomal interactions that require substantially higher forces for disruption. In contrast, low concentrations of low-MW PEG produce only slight compaction with fewer internucleosomal interactions, indicating that nucleosomes remain largely independent under these conditions. These findings provide important insights into how molecular crowding modulates the local chromatin conformation, thereby influencing the accessibility of chromatin to cellular regulatory factors.

## Results

### Force-distance curves of 12-mer polynucleosome

We employed a salt-dialysis method to reconstitute 12 nucleosomes on 11 kbp linear DNA with biotinylated ends. A single polynucleosome construct was tethered between two streptavidin-coated polystyrene beads held in optical traps and transferred to the measurement channel (Figure S1). We used four sizes of PEG (ethylene glycol (EG), PEG400, PEG4000, PEG8000) and tested 5% (w/v), 10% (w/v), and 15% (w/v) for PEG400 and PEG4000 as crowding agents (Table 1). More than 25 independent constant-velocity pulling experiments were conducted under each condition to measure the force-induced disruption of the polynucleosome structure, and both the applied force and bead-to-bead distance were recorded (Figure 1A).

**Table 1.** Crowding agents used in this study.

| **Crowder** | **Size (kDa)** | **Concentration**<br>**(% (w/v))** |
| --- | --- | --- |
| **w/o PEG** | - | - |
| **EG** | 0.06 | 10 |
| **PEG 400** | 0.4 | 5, 10, 15 |
| **PEG 4000** | 4 | 5, 10, 15 |
| **PEG 8000** | 8 | 10 |

**Figure 1.**
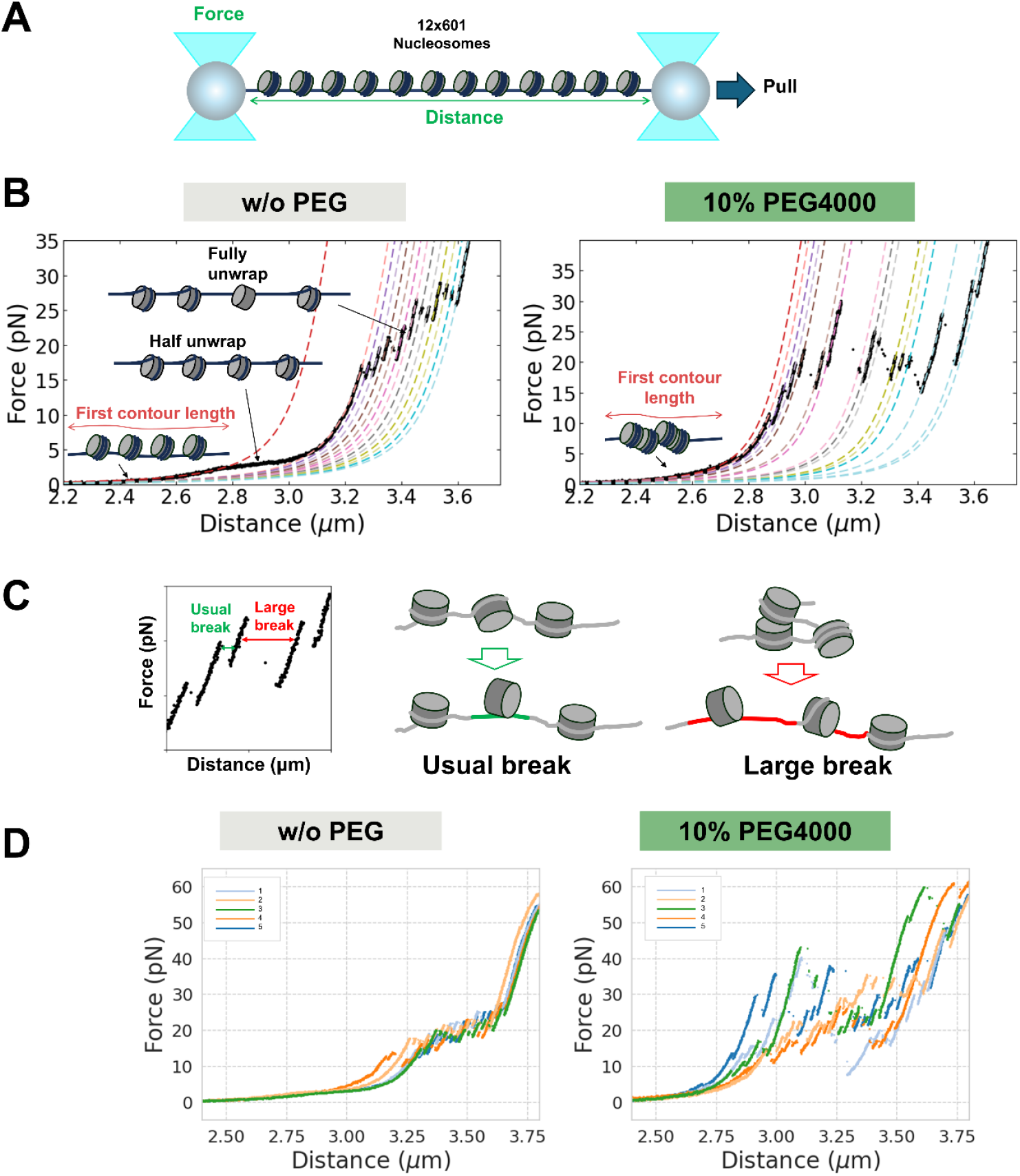
Representative force-distance curves of 12-mer nucleosomal constructs. (A) Schematics of the force-extension experimental setup. (B) Force-distance curve of a 12-mer polynucleosome in the absence of PEG, used as the control, and in the presence of 10% (w/v) PEG4000. In the graphs, black dots represent the measured data, and the dashed lines indicate the fitted WLC curves. DNA unwrapping and polynucleosome structure disruption occur in the pulling experiments. (C) Short and long DNA release events observed in the high-force region. When inner-turn DNA unwrapping occurs, approximately 70 base pairs of DNA are released (usual break). If nucleosomes are associated, longer DNA is released, following the dissociation events (large break). (D) Force-distance curves obtained from five independent pulling experiments in w/o PEG and 10% (w/v) PEG4000.

A representative force-distance curve in the absence of PEG was used as a control and is shown in the left panel of Figure 1B. DNA unwrapping generally occurs in two steps, and these were recorded here. The first step appeared as a gradual force decrease in the low-force region (∼5 pN) at an approximate distance of 2.6 μm, reflecting the unwrapping of outer-turn DNA from all nucleosomes (25). The second step consisted of sharp, sawtooth-like, force drops in the higher force regions (∼20 pN) between 3.2 μm and 3.6 μm. They corresponded to the inner-turn DNA unwrapping around ± 35 bp from the dyad, causing the loss of one nucleosome (25, 26). Model fitting using an extendable wormlike chain (WLC) model provided the length of the construct in each state (dashed lines in Figure 1B). Specifically, the contour length of the first WLC curve (hereafter referred to as the first contour length), shown in red, corresponds to the size of the polynucleosome before structural disruption.

The shape of the force-distance curves reflects the condensation state of the polynucleosome. An example observed with 10% (w/v) PEG4000 is shown in the right panel of Figure 1B. PEG exerted a strong influence in both the low- and high-force regions. The first WLC fitted curve (red dashed line) under 10% (w/v) PEG4000 was shifted to the left compared to that without (w/o) PEG, resulting in a smaller first contour length. This indicated polynucleosome compaction. Large gaps were often observed in high-force regions, and such force drops indicated the release of a loop between the nucleosomes (Figure 1C). Longer DNA release is indicative of the loss of more distal nucleosome-nucleosome associations. A previous study reported a similar loop formation when nucleosomes were bridged with the chromatin protein PRC2 (24). The first contour length and DNA release length in the high-force regions are measures of condensation and nucleosomal associations.

Figure 1D shows representative force-distance curves of five individual constructs from w/o PEG and 10% (w/v) PEG4000 conditions. Detailed analyses of the force-distance curves for all conditions are provided in the following sections.

### Polynucleosome compaction depending on the MW and concentration of PEG

We evaluated the first contour length, which reflects the size of the intact polynucleosome structure (Figure 2). In the absence of PEG, the median first contour length was 3.41 μm. The size of the polynucleosome (12-mer) was estimated to be 0.52 μm (by subtracting the contour length of the 9.3 kbp backbone DNA, which was 2.89 μm). The first contour length decreased with increasing PEG concentration for both PEG400 and PEG4000 (top two panels in Fig. 2). The contour length also decreased with increasing PEG molecular weight, reaching a plateau for PEG4000 (bottom-right panel of Figure 2). The shortest length was observed at 15% (w/v) PEG4000, measuring approximately 3.04 μm, which was a reduction of 0.37 μm compared to the absence of PEG. This corresponded to a 3.5-fold compaction of the polynucleosome region, as the bare DNA was not significantly compacted by PEG (Figure S2). These results demonstrate that high-MW PEG at high concentrations induces substantial compaction of the 12-mer polynucleosomes.

**Figure 2.**
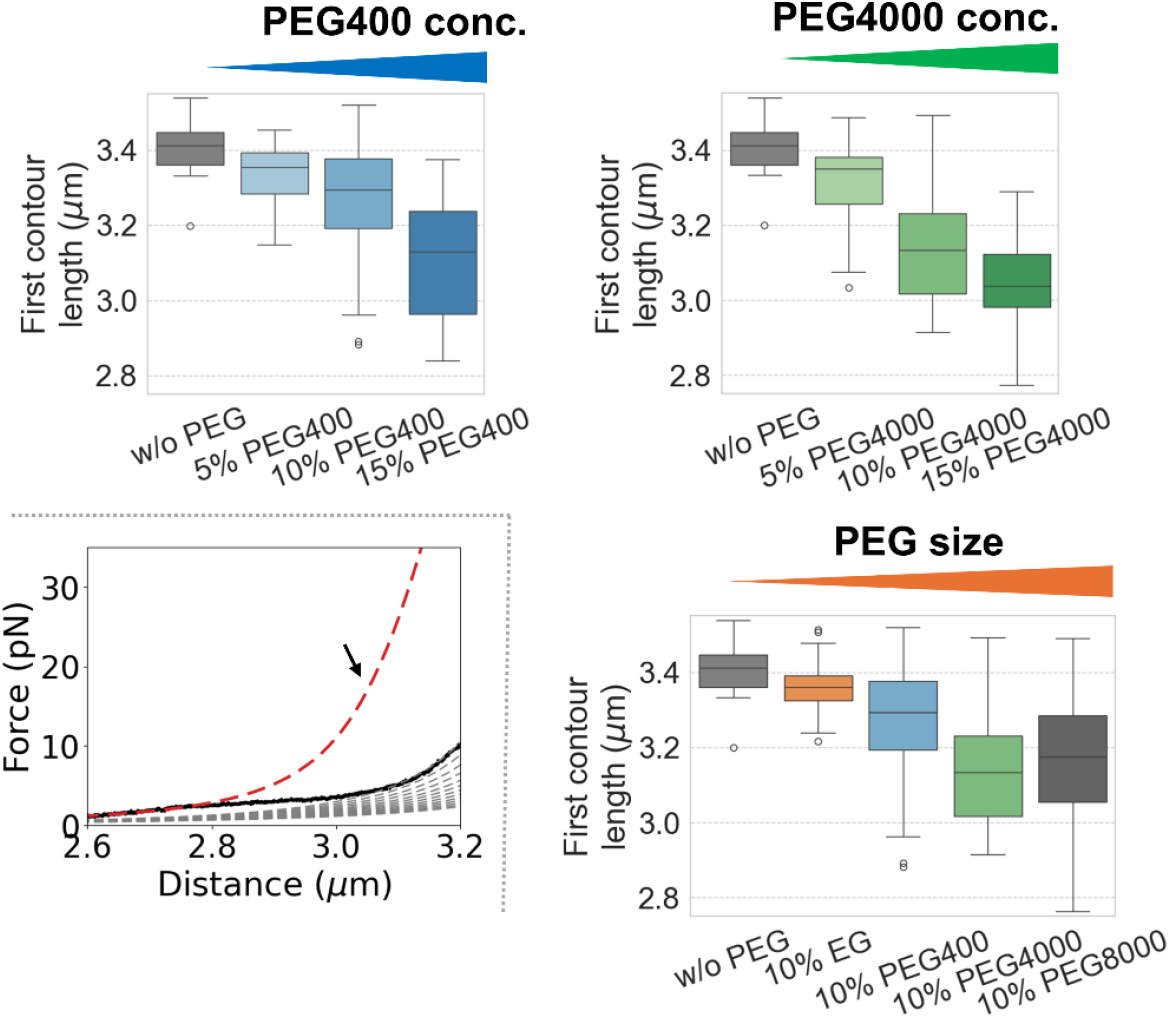
First contour lengths under different conditions. The first contour length is an indicator of the extent of polynucleosome compaction. The figure shows the PEG concentration and size dependent compaction of polynucleosomal constructs.

### Internucleosomal interactions stabilize the compact form of polynucleosomes in the presence of high-MW PEG at high concentrations

To evaluate nucleosome association in the presence of PEG, we analyzed the DNA release lengths in the high-force region. Inner-turn DNA unwrapping from a single nucleosome releases approximately 70 bp, corresponding to the loss of contacts at ±35 bp from the dyad (25, 26). When nucleosomes are associated with each other, the DNA released at each force drop includes both inner-turn DNA unwrapping and simultaneous extension of the DNA (schematic models are shown in Figure S3). Therefore, longer DNA release lengths indicate interactions between the more distal nucleosomes.

We categorized the DNA release lengths into four classes (Figure 3A): Class1 (1–70 bp), corresponding to inner-turn DNA unwrapping (∼70 bp); Class 2 (80–160 bp), representing the single nucleosomal DNA unwrapping (∼146 bp); Class 3 (160–240 bp), associated with disruption of adjacent nucleosome interactions (∼208 bp); and Class 4 (≥ 240 bp), reflecting disruption of more distal internucleosomal associations. In the absence of PEG, 85% of all DNA releases were in Class 1, indicating that most nucleosomes remained separate under these conditions. In contrast, under a high MW and/or concentration (15% (w/v) PEG400, 10% (w/v) and 15% (w/v) PEG4000, and 10% (w/v) PEG8000), Class 3 and Class 4 DNA release was significantly enriched. In the presence of 15% (w/v) PEG4000, where polynucleosomes were most compact (Figure 2), 15% and 18% of force drops were categorized as Class 3 and Class 4, respectively. These results indicated that high-MW PEG at high concentrations enhanced both local and long-range nucleosome associations, promoting polynucleosome compaction.

**Figure 3.**
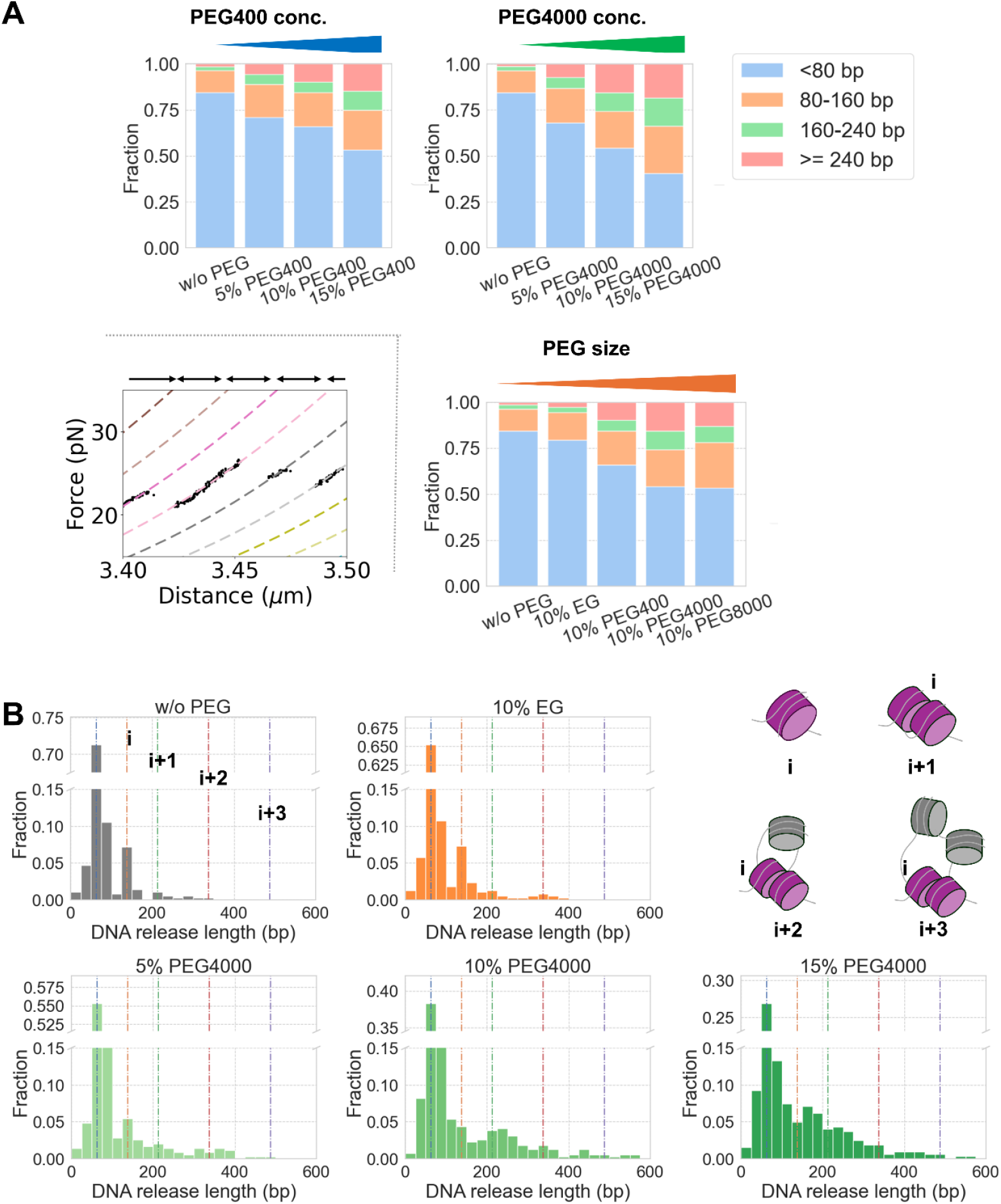
DNA release in the high-force region. (A) PEG concentration and size dependent DNA release lengths. (B) Histograms of released DNA lengths under w/o PEG, 10% (w/v) EG, and 5% (w/v), 10% (w/v), and 15% (w/v) PEG 4000. The middle portion of the y-axis is omitted to better visualize the lower region. Complete histograms for all conditions are presented in Figure S3. Dotted lines indicate estimated release lengths based on nucleosome association schematics shown in Figure S3.

We conducted a detailed analysis of the distribution of DNA release lengths. The histograms in Figure 3B present the results in the absence of PEG and with 10% (w/v) EG, and 5%, 10%, and 15% (w/v) PEG4000 (all histograms are shown in Figure S4). The highest peaks in the histograms consistently appeared at 70 bp across all tested conditions, indicating that strong histone–DNA interactions at approximately ±35 bp from the dyad were maintained both in the absence and presence of PEG. A distinct enrichment at 146 bp, corresponding to the full length of DNA wrapped around a single nucleosome (including both inner and outer turns), was also clearly observed with low-MW PEG at low concentrations (see w/o PEG, 5% (w/v) PEG400, and 10% (w/v) PEG400, as shown in Figure 4 and Figure S4). This suggested that the outer turn occasionally remained wrapped around single nucleosomes, owing to intranucleosomal interactions. Minor enrichments of approximately 200 and 350 bp were also observed, likely corresponding to i-to-i+1 and i-to-i+2 nucleosome interactions under these conditions. Under high-MW PEG at high concentrations (*e*.*g*., 15% (w/v) PEG400, 10% (w/v) and 15% (w/v) PEG4000, and 10% (w/v) PEG8000), similar enrichments at most of these release lengths were observed, although the distributions were broader, and longer DNA releases were preferred. Notably, the 145 bp peak nearly disappeared, particularly at 10% (w/v) and 15% (w/v) PEG4000, indicating that internucleosomal associations are favored over intranucleosomal interactions in such highly compacted states.

**Figure 4.**
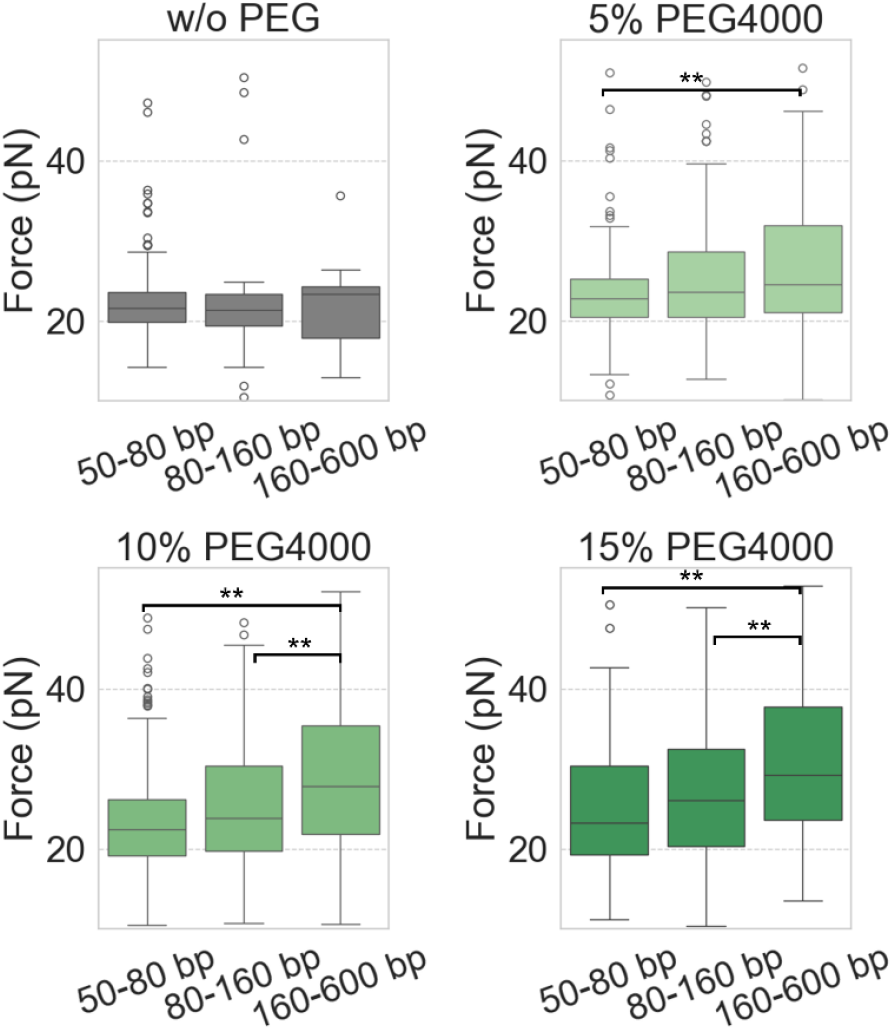
Rupture forces observed in the high-force region in w/o, 5% (w/v), 10% (w/v), and 15% (w/v) PEG 4000, categorized by DNA release lengths: short (50–80 bp), middle (80–160 bp), and long (160-600 bp). The Mann–Whitney U-test was used to determine whether two groups were significantly different: **p ≤ 0.05, *p ≤ 0.1. The graphs for all data are shown in Figure S5. The number of data points for the statistical analyses is shown in Table S2.

### High forces required to break internucleosomal interactions in the presence of high-MW PEG at high concentrations

We evaluated the effect of nucleosome association on the force required for DNA release. Figure 4 shows the rupture forces in the high-force region in the absence of PEG and in the presence of 5, 10, and 15% (w/v) PEG4000 (graphs for all conditions are shown in Figure S5). DNA release lengths were categorized as short (50–80 bp), middle (80–160 bp), and long (160-600 bp) ranges.

In the absence of PEG, no clear difference was observed in the force required to release DNA of different length. In contrast, in the presence of high-MW PEG, the force required to release long DNA (160–600 bp) tended to be higher than that required for short DNA (50–80 bp). Under the most compact conditions with 15% (w/v) PEG4000, a median force of 29.2 pN was required to release long DNA, which was 7.7 pN higher (*i*.*e*., 36% higher) than that required to unwrap inner-turn DNA in the absence of PEG. These results suggest that high-MW PEG at high concentrations stabilizes internucleosomal interactions, strongly restricting DNA exposure to the solvent.

### Mild stabilization of inner-turn DNA-histone interactions in individual nucleosomes in the presence of PEG

Finally, we analyzed the unwrapping forces and kinetics of nucleosomes to evaluate whether PEG affects histone–DNA interactions within individual nucleosomes.

First, we examined the unwrapping forces generated by 12-mer nucleosome arrays. To minimize the influence of nucleosome associations, we focused only on DNA release lengths of 60–80 bp under conditions where no events showed DNA release exceeding 200 bp (the detailed procedure is described in the Materials & Methods). The results are shown in Figure 5A. The median unwrapping force increased by 1.0–1.4 pN in the presence of PEG compared to that in the absence of PEG.

**Figure 5.**
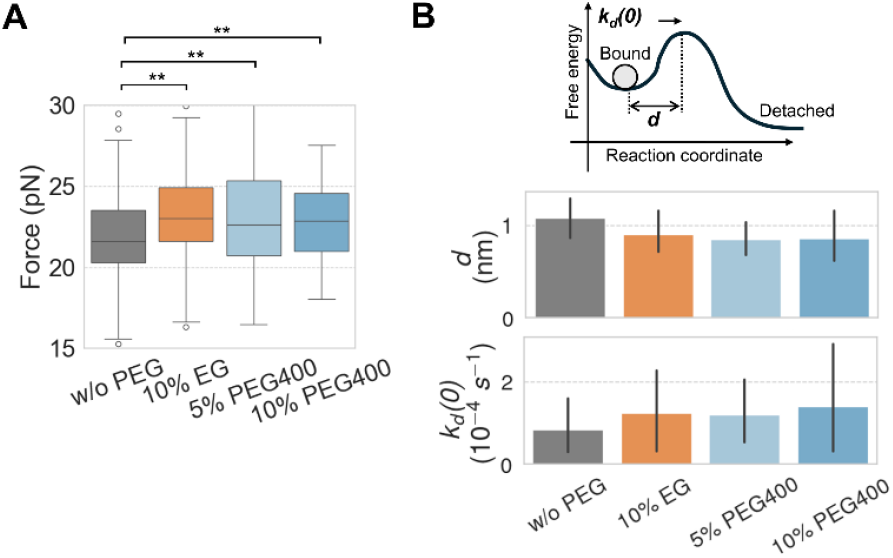
(A) DNA unwrapping force required for the DNA release length of 60–80 bps, representing the force required to remove the inner-turn DNA without nucleosomal association. The Mann–Whitney U-test was used to determine whether two groups were significantly different: **p ≤ 0.05, *p ≤ 0.1. (B) Kinetic parameters for the disruption. The error bar indicates bootstrap confidence intervals of 90%. There are no significant differences in the parameters. The number of data points for the statistical analyses is shown in Table S3.

To assess the effect of PEG on the disruption kinetics, we calculated the kinetic parameters, the zero-force velocity (*k*_*D*_(0)), and the distance from the ground state to the transition state (*d*) (Figure 5B) for the 12-mer array using a previously described method (25). No significant changes were observed in the kinetics under the different PEG conditions. Therefore, we concluded that the increased force observed in the presence of PEG resulted from increased binding stability between histones and DNA, rather than from changes in kinetics.

In summary, our data suggest that histone–DNA interactions corresponding to the inner-turn DNA region (70 bp DNA release, ±35 bp from the dyad) are stabilized in the presence of PEG, even at low PEG concentrations where internucleosomal interactions are barely observed.

## Discussion

Our study provides insights into how molecular crowding influences polynucleosome structures at the molecular level. First, PEG promoted polynucleosome compaction in a size- and concentration-dependent manner. Second, internucleosomal interactions were markedly stabilized with high-MW PEG at high concentrations, whereas with low-MW PEG at low concentrations, nucleosomes remained largely separated, although DNA unwrapping was slightly inhibited. Third, a substantial mechanical force (∼30 pN) was required to disrupt these internucleosomal interactions and release DNA from highly compacted polynucleosomal structures under high-MW PEG at high concentrations.

### Compaction occurs in the polynucleosome region, not in the DNA region

Our construct consisting of 8.3 kbp of bare DNA and a 2.5 kbp polynucleosome region, formed a compact structure in the presence of PEG (Figure 2).

Both the DNA and polynucleosomes are known to undergo condensation in the presence of PEG. Circular dichroism measurements of calf thymus DNA revealed a critical PEG concentration threshold for DNA condensation, occurring at approximately 20% (w/v) PEG2000–6000 in 100 mM NaCl (27). Furthermore, single-molecule fluorescence microscopy consistently showed that T4 DNA did not condense until 20% (w/v) PEG10000 (16). In contrast, magnetic tweezers experiments (20) showed that 46 kbp lambda DNA forms a globular structure in the presence of 13–22% (w/v) PEG1500. This structure unfolds under very low force (0.3–1.4 pN). However, our tweezers experiments did not show a similar increase in force for the 11 kbp bare DNA used for polynucleosome reconstitution (Figure S2). This may be because our DNA was four times shorter than lambda DNA, resulting in a much smaller force increase that was undetectable in our experimental setup. As such, we concluded that the compaction observed in our force-distance curves reflects the structural changes within the polynucleosome region in the presence of PEG.

Few studies have been conducted on polynucleosome condensation under crowded conditions. Fluorescence microscopy studies have shown that polynucleosomes reconstituted on 169 kbp T4 DNA (at various nucleosome densities ranging from half to fully reconstituted arrays) undergo compaction with increasing concentrations of both PEG10000 and BSA (16, 17). Notably, they reported no critical concentration for polynucleosome condensation, in contrast to that observed for the DNA condensation. Our data also showed slight compaction at 5% (w/v) PEG400 and 5% (w/v) PEG4000, with the polynucleosome size gradually decreasing with an increase in the PEG concentration (Figure 2, upper panels). These data suggested that the increased depletion forces arising from the excluded volume effect of crowder molecules, which become stronger at higher concentrations, progressively promote polynucleosome compaction. Although our construct contained a much smaller polynucleosome region (2.5 kbp with 12 nucleosomes) than the 169 kbp T4 DNA used in previous studies, the results exhibited similar crowder concentration-dependent compaction. Our data thus provide evidence that such compaction can occur even in small polynucleosome regions.

### PEG-size dependent compaction through excluded volume effect screening

Here, we report that the degree of polynucleosome compaction depends on the molecular weight of PEG, with higher-MW PEG molecules causing greater polynucleosome compaction.

Theoretical studies have suggested that the excluded volume effect is determined not only by the crowder’s occupancy but also by the geometry and size (28). A notable observation was reported regarding the compactness of intrinsically disordered proteins (IDPs) of approximately 50 amino acids, investigated using single-molecule FRET in the presence of PEG as a crowder (29). The study showed that the IDPs became more compact not only with increasing concentration, but also with increasing PEG size. The compaction of the IDPs plateaued at the size of PEG4600. We also observed similar PEG-size-dependent compaction, and it reached a plateau at a certain PEG size (Figure 2, bottom right panel). A renormalized Flory-Huggins model that accounts for the interpenetration of polymer chains was proposed to explain the PEG-size dependency of IDP compaction (29). This phenomenon is referred to as excluded volume effect screening, because the interpenetration of polymer chains reduces the excluded volume effect that would otherwise be observed with spherical crowders of the same size. The similarity in the PEG-size dependency between IDPs and polynucleosomes suggests that excluding volume effect screening through the interpenetration of PEG chains, both within PEG molecules and between PEG and polynucleosomes, contributed to compaction in our system. This is particularly plausible, given that histone tail domains that extend from the nucleosome core are intrinsically disordered and could facilitate such interpenetration with PEG chains.

### DNA exposure is strongly restrained by nucleosomal associations at high-MW PEG and high concentrations

Our analysis of the force-induced disruptions in the high-force region reveals significant enrichment of DNA releases of 160–240 bp and ≥ 240 bp at high-MW PEG and high concentrations (Figure 3A), indicating that stable internucleosomal interactions contribute to the pronounced compaction of the polynucleosome. As shown in Figure 4, approximately 30 pN of force was required to disrupt the interactions; this amount of force is 36% higher than that required for typical DNA unwrapping in the absence of PEG.

Herein, we discuss the potential causes of these nucleosome associations. Previous studies have shown that histone tails contribute to the formation of higher-order chromatin structures (30). In particular, the N-terminal tails of H4 and H3 are involved in both intra- and internucleosomal interactions, promoting the compaction of nucleosomal arrays (31, 32). The tails primarily interact with DNA (33, 34). In addition, the basic patch of the H4 tail interacts with the H2A/H2B acidic patch, both within the nucleosome and between adjacent nucleosomes (35). These highly dynamic and heterogeneous interactions (36, 37) are likely enhanced by the depletion forces under crowded conditions. The fuzzy nature of these interactions likely contributed to the broad distribution of release lengths in the compacted forms (Figure 3B and S4).

In addition to tail-mediated interactions, nucleosome entanglement, as mentioned in a previous work (38), may be facilitated under crowded conditions. A simulation study suggested that the self-entanglement of linear polymers is prevalent in crowded environments (39). Although our system differs from simple linear polymers, it is likely that nucleosome entanglement occurred under our experimental conditions. The prolonged DNA release observed in our experiments is therefore likely attributable to both histone tail-mediated interactions and nucleosome entanglement.

Finally, we propose a model for the structure of polynucleosomes under varying PEG concentrations and sizes, as illustrated in Figure 6. Under low PEG concentrations with low-MW PEG, weak depletion forces maintain nucleosomes in a largely separated state, whereas DNA-histone interactions become slightly stabilized (Figure 5A). In this regime, occasional histone tail-mediated interactions or an entanglement in a single nucleosome occurred, corresponding to 147 bp release in the pulling experiment (*e*.*g*., 10% (w/v) EG in Figure 3B), but internucleosomal interactions were rarely observed. Conversely, high PEG concentrations combined with high-MW PEG generated strong depletion forces that pushed nucleosomes toward each other, promoting histone tail bridging and multinucleosome entanglement and thereby driving polynucleosome compaction (*e*.*g*., 10% (w/v) and 15% (w/v) PEG4000 in Figure 3B). In these highly compact conformations, DNA accessibility to the polynucleosome structure was strongly suppressed (Figure 4).

**Figure 6.**
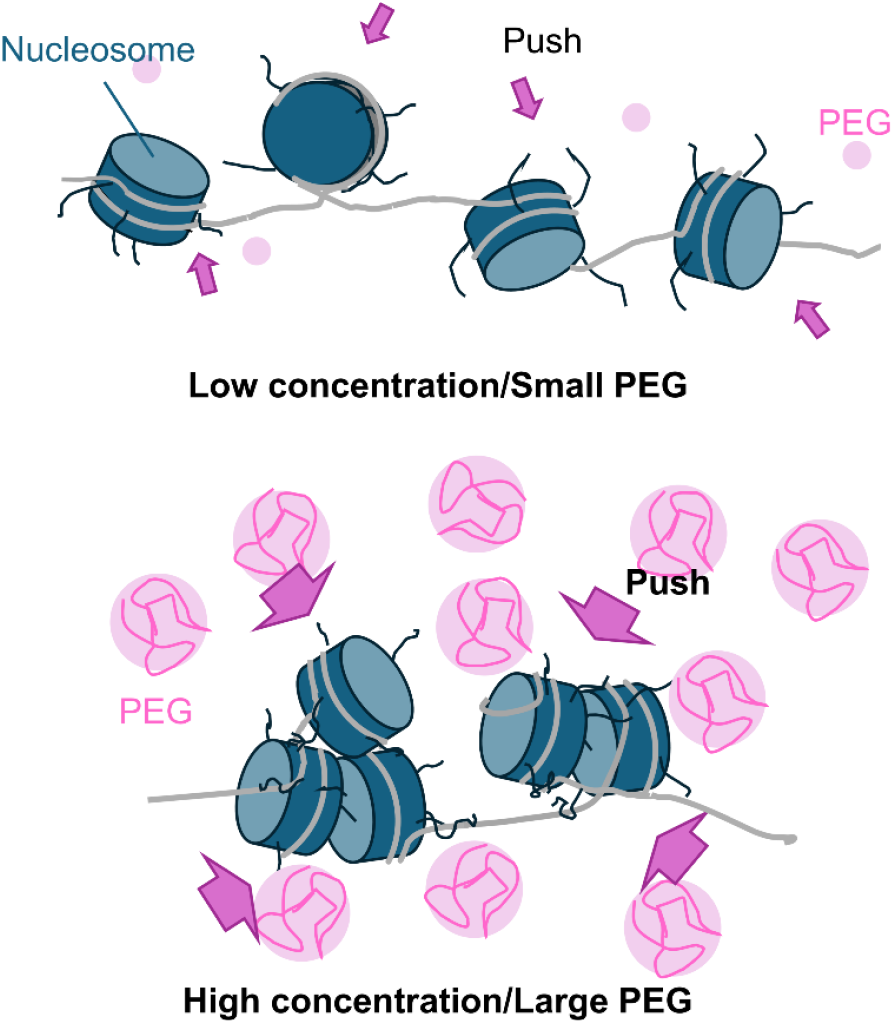
Proposed structural models of polynucleosome under different crowded conditions. Under low concentrations and small PEG, nucleosomes remain separated from each other, and slight stabilization of DNA-histone interactions occurs within individual nucleosomes. In contrast, under high concentrations and larger PEG molecules, nucleosomes are strongly pushed by depletion forces to interact with one another, forming a compact structure in which DNA becomes less accessible.

### Biological relevance

Chromatin condensation is the key regulator of transcription and replication (1, 2). Based on our findings, stable internucleosomal interactions formed under highly crowded conditions likely hinder the access of regulatory proteins, such as epigenetic reader/writer proteins, chromatin remodelers, and polymerases. Notably, the RNA polymerase from *E*.*coli* has been reported to stall at a force of approximately 25 pN (40), although similar data for eukaryotic RNA polymerases have not yet been reported. This force is lower than that required to disrupt inter-nucleosome associations with high-MW PEG at high concentrations, but higher than the force needed to unwrap DNA with low-MW PEG at low concentrations. These observations suggest that high-molecular crowding slows the passage of motor proteins, such as RNA polymerase, by enhancing nucleosome-nucleosome interactions.

The chromatin-binding protein PRC2 has been reported to enhance internucleosomal interactions to a level similar to that observed in the present study (24). Our findings demonstrate that the excluded volume effect due to molecular crowding alone, without the involvement of specific chromatin-binding proteins, can induce internucleosomal interactions of comparable strength. Molecular crowding may also contribute to the establishment of specific chromatin states, such as the mitotic chromosome structure. Indeed, Iida *et al*. reported that the molecular density surrounding mitotic chromosomes increased during mitosis (11). This observation suggests that living systems may possess mechanisms that precisely regulate chromatin structure through the spatial and temporal modulation of molecular crowding, thereby controlling DNA accessibility for transcription and replication.

## Materials and Methods

### Preparation of DNA

The plasmid containing 12 repeats of the 147 bp Widom601 (41) (12×601) sequence with a 61 bp-linker was assembled into 8.3 kbp pKYB1 (NEB) using the Gibson Assembly method (Thermo Fisher Scientific). The protocol for DNA preparation has been previously described (42). The final plasmid (11.0 kbp) was transformed to *E*.*coli* cells, and the amplified DNA was purified using a plasmid mini-prep kit (Nippon Genetics). The DNA was then linearized using *Eco*R I digestion (Takara Bio). Both ends of the DNA were biotinylated with Biotin-14-dATP (Thermo Fisher Scientific) using the Klenow fragment (3’→5’ exo^-^) (NEB) at 37 °C for 30 mins. The resulting biotinylated DNA was purified using a gel-PCR extraction kit (Nippon Genetics).

Additionally, 147 bp DNA was prepared as a competitor for nucleosome reconstitution (43). This fragment was amplified from the pUC19 plasmid by PCR using the KOD-Plus-high-fidelity PCR enzyme (Toyobo). Purification was accomplished by non-denaturing polyacrylamide gel electrophoresis using a Prep Cell apparatus (Bio-Rad).

### Preparation of histone octamer

We prepared an octamer consisting of H2A, H2B, an H3K9me3-mimic, and H4, as described previously (44, 45). Briefly, His-tagged H2A, H2B, H3.2 K9C/C110A, and H4 were expressed in *E*.*coli* and purified using Ni-NTA agarose beads (Qiagen). His-tag was removed by thrombin protease cleavage. Proteins were further purified using Mono S columns (Cytiva). To introduce tri-methylation on K9 of H3, 5 mg/ml H3 in alkylation buffer containing 1 M HEPES-NaOH (pH 7.8), 4 M guanidinium chloride 10 mM D/L-methionine, and 20 mM DTT were incubated with a 20-fold excess (by weight) of (2-bromoethyl) trimethylammonium bromide (Sigma-Aldrich) at 50 °C for 150 min, followed by the addition of DTT to a final concentration of 10 mM and incubation at 50 °C for 150 min. The reaction was stopped by the addition of 2-mercaptoethanol to a final concentration of 880 mM. The resulting H3K9me3 was desalted on a PD-10 column (Cytiva). All the purified histone proteins were lyophilized. H2A, H2B, H3K9me3-mimic, and H4 proteins were mixed in denaturing buffer containing 50 mM Tris-HCl (pH 7.5), 7 M guanidine-HCl, and 20 mM 2-mercaptoethanol. The mixture was dialyzed against a refolding buffer containing 2 M NaCl in 10 mM Tris-HCl (pH 7.5) and 2 mM 2-mercaptoethanol to reconstitute the histone octamers. The octamer was purified using Superdex 200 gel-filtration column chromatography (Cytiva).

### Nucleosome reconstitution

Nucleosomes were reconstituted in 12 × 601 DNA, as previously described (42, 43). Briefly, we mixed 60 ng/μl biotinylated 12 × 601 DNA with an equal amount of competitor DNA and histone octamers. The salt concentration was adjusted to 2 M. The concentration gradually decreased to 250 mM over 33 h of dialysis. The buffer was replaced with 10 mM Tris-HCl (pH 7.5) by further dialysis. The reconstituted polynucleosome was kept at -80 °C, after adding glycerol to a final concentration of 5% (v/v).

### Optical tweezers force-extension experiment

We used a dual-trap optical tweezer system, mTrap (Lumicks), to perform DNA pulling experiments at a constant velocity. The flow-cell configuration is shown in Figure S1. The streptavidin-coated polystyrene beads, with a diameter of 1.76 μm (Spherotech), were captured by the optical traps. A biotinylated polynucleosomal DNA construct was tethered between the beads in the running buffer: 20 mM HEPES-NaOH, pH 7.5, 100 mM NaCl, 0.5 mM EDTA, 0.02% (v/v) Tween 20, and 0.2% (w/v) BSA. The DNA tethered beads were then transferred into a measurement buffer: 20 mM HEPES-NaOH, pH 7.5, 100 mM NaCl, 0.5 mM EDTA, 0.02% (v/v) Tween 20, 0.2% (w/v) BSA, and different concentration of PEG, where it was stretched approximately from 2 to 4 μm at a constant velocity of 20 nm/s until the DNA was fully unwrapped from the histone core proteins.

The mTrap system was equipped with a position-sensitive detector to measure bead displacement. The force applied to the bead (*F*) is related to the displacement (*x*) by the trap stiffness (*κ*_*f*_) as,

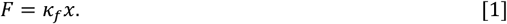

K_f_ was determined by analyzing the power spectrum of the Brownian motion of a trapped bead in the measurement buffer. The power spectral density *S*(*f*) at frequency *f* is given by

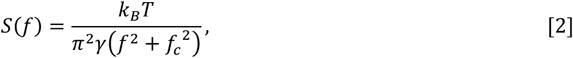

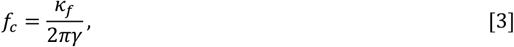

where *k*_*B*_ is the Boltzmann constant, *T* is temperature, and *γ* is the drag coefficient(46). *γ* is 3*πηd* for a spherical bead (*d* is the bead diameter, and *η* is the dynamic viscosity of the fluid). *γ* was obtained from an independent force calibration as described in the literature (47), by oscillating the piezo stage at 17 Hz in the respective measurement buffer. *η* derived from *γ* is shown in Table S1.

### Calculation of DNA release length

The pulling experiment for the polynucleosome shows the stepwise release of DNA in the low (∼5 pN) and high (≥10 pN) force regions. The force-distance curve consists of discontinuous segments separated by gradual or sudden force drops in the low- and high-force regions, respectively. We extracted force drops with a decrease greater than 0.2 pN and an extension of 50 nm or more. These segments were selected from 0-10 and 10-50 pN for the low-and high-force regions, respectively. Each segment was fitted globally using the extendable WLC model proposed by Odijk (48),

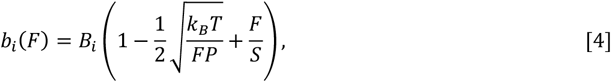

where *b*_*i*_ (*F*) is the observed distance between the beads of *i* -th segment at force *F* ; *B*_*i*_ is the contour length of the segment; and *P* and *S* are the persistent length and stretch modulus, respectively, fitted as common parameters across all segments. The length of DNA released during each event was calculated as the difference between the two contour lengths. Values are presented in base pairs (bp), assuming that one base pair corresponds to 0.34 nm.

### Calculation of kinetic parameters for inner-turn DNA unwrapping

The disruption kinetics of the inner turns of individual nucleosomes in the 12-mer array were calculated as previously described (25). When *N* nucleosomes remain on the array, the average force *F*^∗^ required to disrupt a single nucleosome is expressed as

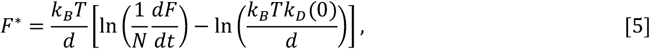

where *k*_*D*_(0) is the rate constant for disruption in the absence of an external force, and *d* is the distance between the bound state and the activation barrier peak. *N* for each segment was calculated from the difference between the contour length of the last segment and that of the respective segment. The loading rate *dF*/*dt* was set to 0.4 pN/s. To minimize the influence of nucleosomal association, only molecules without long DNA release events (>200 bp) were included when calculating the kinetic parameters. Conditions were considered only when at least ten data points remained. Force drops corresponding to a DNA release of 60–80 bp were used to calculate *F*^∗^, focusing specifically on inner-turn DNA unwrapping.

All the analyses described above were performed with Pylake Python library (49).

## Acknowledgments

We appreciate our valuable discussions with W. S. Chan, H. Ishida, A. Matsumoto, J. Ono, and S. Sakuraba. We thank J. Kato for her assistance in preparing polynucleosomes. We are also grateful to L. Chaubet for his guidance on the force measurements using mTrap (dual-trap optical tweezers). This work was supported by JSPS KAKENHI (grant numbers JP18H05534 [H Kurumizaka, H. Kono], and JP22K06176, JP23K05726, and JP24H00884 [T.S.], JP23H05475 and JP24H02328 [H. Kurumizaka], and JP22K06179 [S.S.]), QST Strategic President’s Fund “Budding Research” [A.K.], BINDS from AMED (grant numbers JP25ama121025 and JP25ama121009) [H. Kono, H. Kurumizaka], and Japan Science and Technology Agency CREST (grant number JPMJCR24T3 [H. Kurumizaka]).

## Author Contributions

T.S. and H. Kono: Conceptualization. T. S., S. S., and H. Kurumizaka: sample preparation. T. S. and Y. H.: data collection; T. S.: formal analysis. T. S., A. K., and H. Kono wrote the original drafts and edited the manuscript. All authors have reviewed and approved the final manuscript.

## Data Availability

Data will be made available on request to T.S. and H. Kono.

## Notes

### Competing Interest Statement

The authors have declared no competing interest.

