## Supplemental Figures S1 to S5 and Tables S1 to S3 for "Molecular Crowding-Driven Nucleosome Interactions Revealed Through Single-Molecule Optical Tweezers"

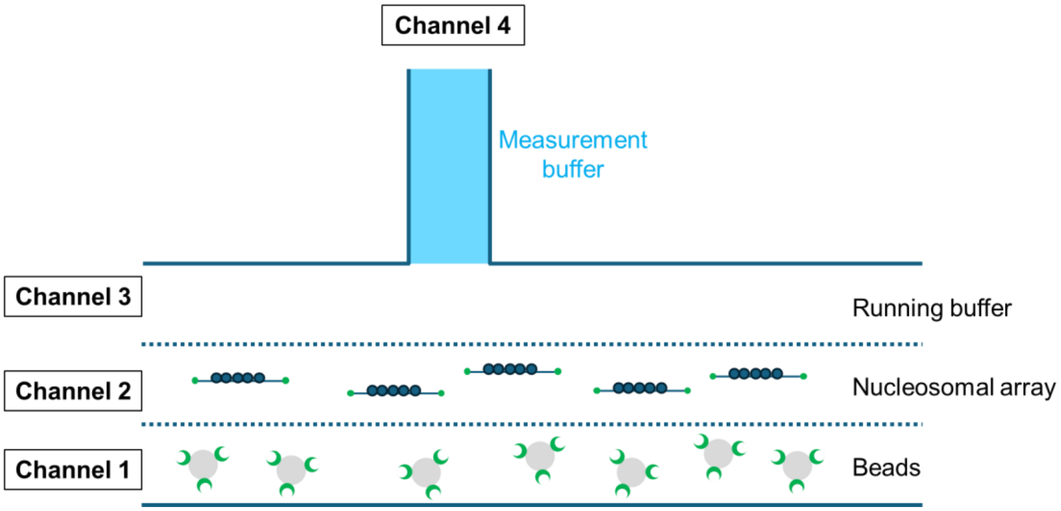


**Fig. S1.** Flow cell configuration. For each pulling experiment, two streptavidin-coated beads were trapped in Channel 1 and moved to Channel 2 to allow tethering of a nucleosomal construct between them. Once the nucleosomal construct was loaded, it was transferred to Channel 4, where the pulling experiments were performed.

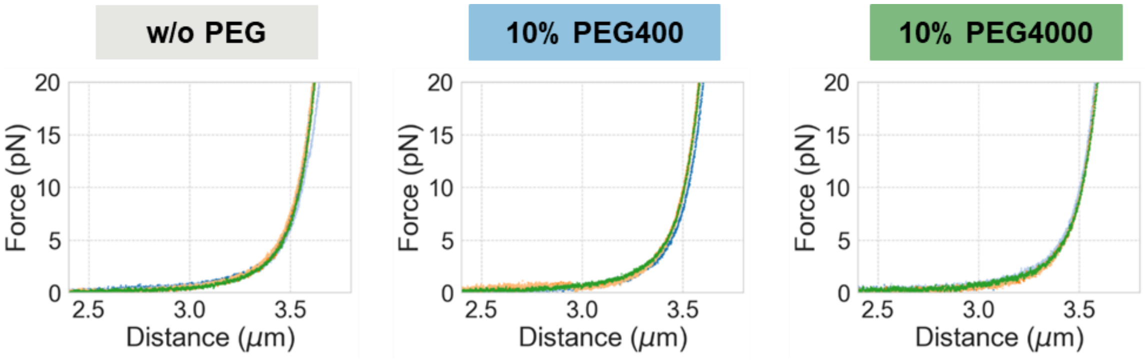


| | Contour length (μm) | Persistence length (nm) | Stretching modulus (pN) |
|---|---|---|---|
| **w/o PEG** | 3.75 ± 0.04 | 56.5 ± 9.0 | 1414±158 |
| **10% PEG 400** | 3.73 ± 0.01 | 52.9 ± 5.4 | 1425±120 |
| **10% PEG 4000** | 3.72 ± 0.05 | 50.3 ± 5.4 | 1446±134 |

**Fig. S2.** Five representative force-extension curves of 11 kbp bare DNA in the presence and absence of PEG and the obtained mechanical parameters. Data are shown as mean ± STD.

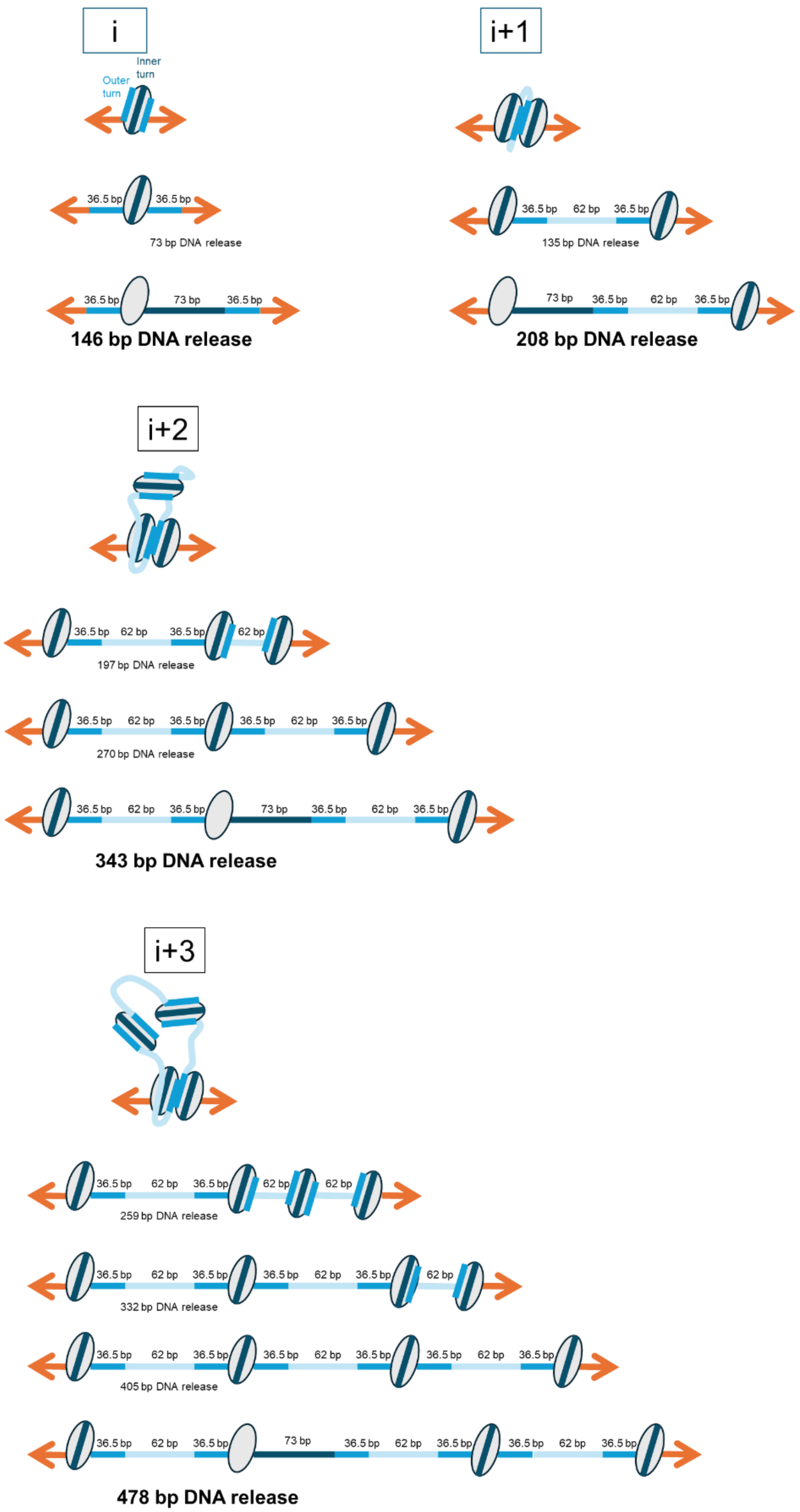


**Fig. S3.** Schematic explanation of DNA release lengths following nucleosomal dissociation events observed at force rupture points in the force-extension curve. DNA release events involving interactions with nucleosomes at positions i, i+1, i+2, and i+3 are illustrated.

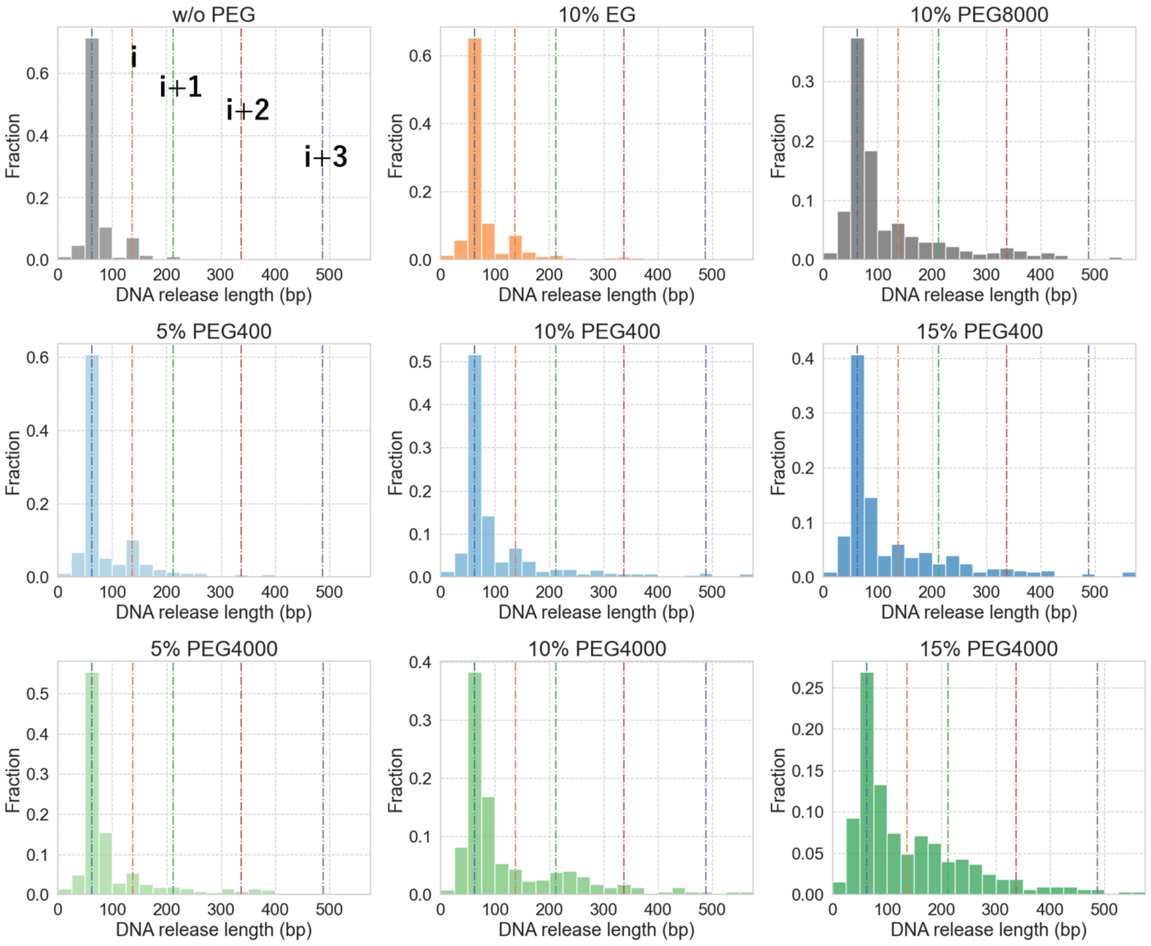


**Fig. S4.** Number of DNA fragments released in the high-force regions. This figure includes all data collected in the study.

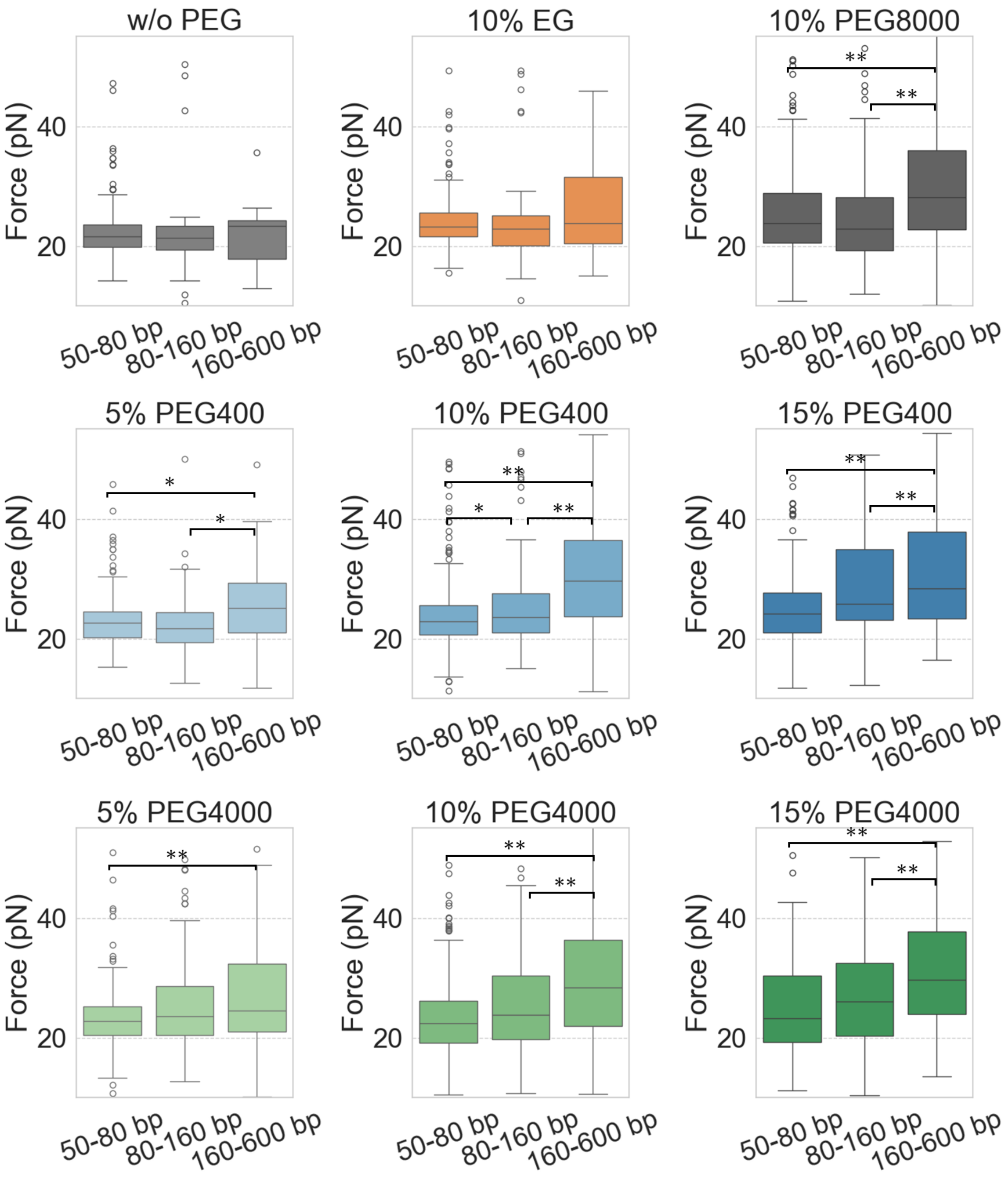


**Fig. S5.** Rupture forces were observed in the high-force region and categorized by DNA release length: short (50–80 bp), medium (80–160 bp), and long (160-600 bp). The Mann–Whitney U-test was used to determine whether the two groups were significantly different: **p ≤ 0.05, *p ≤ 0.1. This figure includes all the data collected in this study. The number of data points used for the statistical analyses is shown in Table S2.

**Table S1.** Dynamic viscosity obtained from active calibration.

| | **Dynamic viscosity (mPa・s)*** |
|---|---|
| **w/o PEG** | 0.982 ± 0.023 |
| **10% EG** | 1.221 ± 0.032 |
| **5% PEG 400** | 1.189 ± 0.012 |
| **10% PEG 400** | 1.426 ± 0.017 |
| **15% PEG 400** | 1.631 ± 0.057 |
| **5% PEG 4000** | 1.627 ± 0.028 |
| **10% PEG 4000** | 2.562 ± 0.040 |
| **15% PEG 4000** | 4.145 ± 0.230 |
| **10% PEG 8000** | 3.787 ± 0.069 |

* Data are shown as mean ± SD

**Table S2.** Number of data points in Figures 4 and S5.

| Condition | DNA release size | Number of data |
|---|---|---|
| **w/o PEG** | 50–80 bp | 285 |
| | 80–160 bp | 43 |
| | 160–600 bp | 13 |
| **10% EG** | 50–80 bp | 276 |
| | 80–160 bp | 58 |
| | 160–600 bp | 21 |
| **5% PEG400** | 50–80 bp | 179 |
| | 80–160 bp | 51 |
| | 160–600 bp | 32 |
| **10% PEG400** | 50–80 bp | 255 |
| | 80–160 bp | 80 |
| | 160–600 bp | 67 |
| **15% PEG400** | 50–80 bp | 150 |
| | 80–160 bp | 72 |
| | 160–600 bp | 84 |
| **5% PEG4000** | 50–80 bp | 218 |
| | 80–160 bp | 65 |
| | 160–600 bp | 47 |
| **10% PEG4000** | 50–80 bp | 179 |
| | 80–160 bp | 79 |
| | 160–600 bp | 102 |
| **15% PEG4000** | 50–80 bp | 99 |
| | 80–160 bp | 86 |
| | 160–600 bp | 112 |
| **10% PEG8000** | 50–80 bp | 177 |
| | 80–160 bp | 102 |
| | 160–600 bp | 90 |

**Table S3.** Number of data points in Figure 5.

| Condition | Number of Data |
|---|---|
| **w/o EG** | 195 |
| **10% EG** | 180 |
| **5% PEG400** | 86 |
| **10% PEG400** | 79 |
